# Identification of specific lipid-protein interactions in dividing cells using lipid-trap mass spectrometry

**DOI:** 10.1101/2024.12.13.627510

**Authors:** Andrea Paquola, Cagakan Ozbalci, Elisabeth M. Storck, Stephen J. Terry, Clare E. Benson, Ulrike S. Eggert

## Abstract

Cells actively maintain complex lipidomes that encompass thousands of lipids, however, many of the roles of these lipids remain unexplored. Specific interactions between lipids and membrane proteins are a likely reason for the evolutionary conservation of complex lipidomes. We report the development of a technique, named lipid-trap mass spectrometry (LTMS), to systematically study protein-lipid interactions directly captured from mammalian cells. LTMS uses immunoprecipitation of GFP-tagged proteins expressed in HeLa, followed by lipidomic analysis of lipids bound to the GFP-tagged protein. We applied LTMS to cell division to illustrate the technique. We chose this process because membranes regulate their lipid composition as they undergo major changes during cytokinesis and many cytokinetic proteins, including RACGAP1 and ESCRTIII components CHMP4B and CHMP2A, are membrane-associated. Using LTMS, we found that RACGAP1 and CHMP4B associate with specific lipid species in dividing compared to non-dividing cells. We expand our understanding of lipid diversity during cell division and present a general approach to explore lipid-protein interactions to further our understanding of the roles of lipids in mammalian cells.

## Introduction

Lipids are important biological molecules that participate in many essential cellular processes and primarily reside in the plasma membrane and the membranes of internal organelles. Mammalian cells use evolutionarily conserved pathways to produce thousands of chemically distinct lipids, which requires substantial investments of energy and resources. Phospholipid diversity is defined by variations in polar head groups and hydrophobic side chains. Lipid families are commonly named according to their head groups (e.g. serine). Side chains can vary in length and the number of double bonds they contain, which is also reflected in the lipid’s name, e.g. PS 18:1_18:1 is a phosphatidyl serine (PS) with 2 side chains that are both 18 carbons long and have 1 double bond. In the past, it was thought that lipids’ principal role was to form water-impermeable barriers to compartmentalize different cellular reactions ^1^. It is now clear that lipids also have many other functions, for example, they play key signaling roles and are necessary in metabolism ^1–3^. However, none of these roles explain why cells produce (and therefore presumably require) such large and diverse lipidomes, especially since much of the chemical diversity lies in the hydrophobic lipid side chains that are normally buried within lipid bilayers. Emerging literature suggests that specific lipids can interact with membrane proteins ^4, 5^ and can be required for the functions of these proteins, raising the possibility that specific and regulated lipid-protein interactions could be a reason for the need for lipid diversity. However, so far systematic studies of lipid-protein interactions in mammalian cells have been limited, both due to a lack of general understanding of lipid regulation and because there were limited techniques to do so.

It is becoming increasingly clear that lipids are essential for protein activities ^5–7^. The lipid environment can affect the binding of agonists and antagonists to GPCRs ^7^. Lipids also play key roles in the folding and oligomerization state of membrane-associated proteins ^4, 6, 8^. For example, the addition of cardiolipin to the sugar transporter SemiSWEET was shown to cause a change in the equilibrium from monomer to dimer ^4^. In addition, experiments in membrane models showed that hydrophobic mismatches due to a difference between the hydrophobic thickness of a membrane and a transmembrane protein segment could induce protein clustering, leading to changes in both their conformation and function ^9, 10^. The study of lipid-protein interactions and how they modulate protein activity is of particular interest for pharmaceutical development, considering that up to 60% of drug targets are located in the plasma membrane ^11^. Different techniques have been developed to investigate lipid-protein interactions *in vitro*, for example, lipid overlay and lipid pull-down assays. In these assays, lipids are immobilized on a solid support. Proteins are added to the immobilized lipids and bound proteins are analyzed via immunodetection. While this technique has resulted in valuable information, its main downside is that lipid-protein interactions are studied under artificial conditions. Chemically derivatized lipids have also been used to study lipid-protein interactions ^12, 13^. These lipids are often characterized by a bifunctional unit such as a photo-activatable group to allow cross-linking with the protein, and an azide or alkyne group that can be “clicked” with a tag, such as biotin, and isolated through affinity purification, with subsequent identification of protein interactors by mass spectrometry. This technique can be very informative but requires complex and highly specialized chemical synthesis.

Of the techniques developed to study lipid-protein interactions in cellular assays, affinity purification lipidomics using immunoglobulin-based pulldowns is the most straightforward. This method was applied to enzymes involved in the ergosterol biosynthesis pathway in yeasts. A variety of small metabolites, including lanosterol and ergosterol, were isolated ^14^. Native protein mass spectrometry is another promising method that has led to recent successful insights ^4, 5, 15, 16^. The rationale for the mass spectral analysis of native protein assemblies is based on the capacity of mass spectrometry to measure small changes in the mass-to-charge ratio (*m/z)*, which can indicate the binding of small molecules, including lipids ^17^. This technique has already given a great contribution to our understanding of lipid-protein interactions, especially in bacteria. However, we have lacked a technique to systematically study membrane-associated proteins in mammalian cells, yet this is imperative to better understand lipid diversity and shed light on how lipids are dysregulated in disease.

Lipids can associate with membrane-bound proteins in different ways, classified as annular, non-annular or bulk lipid binding. Non-annular lipids bind to membrane-associated proteins directly, often in specific binding pockets, and have restricted mobility. Annular lipids form a “ring”, i.e. annulus, around the protein and are in dynamic exchange with the surrounding bulk lipids ^18^. Annular lipids play crucial roles in maintaining the amphipathic environment necessary for the folded state of proteins ^19^. Bulk lipids might also be crucial for protein activity, for example, by controlling their localization or defining the local lipid environment such as microdomains or areas of specific curvature. Understanding how membrane-associated proteins interact with their surrounding lipids, both tightly bound non-annular and more dynamic halo-like annular or bulk lipids is essential to fully investigate their many biological functions. We report here the development of a technique that allows the isolation and identification of lipids associated with tagged proteins extracted from their native environment in live cells. We name this technique lipid-trap mass spectrometry (LTMS) and apply it in a model study of membrane-associated proteins during cytokinesis, the final stage of cell division.

When cells divide, the plasma membrane and all internal organelles, including the lipids they are composed of, are rearranged and distributed amongst daughter cells ^20, 21^. The field has a good understanding of which proteins are involved in cell division, how they are regulated and which tasks they perform. However, much less is known about the lipids and the membranes they reside in. We reported that cells regulate their lipid composition with high temporal and spatial precision as they divide. Cells specifically accumulate certain lipids in the midbody, the small structure that forms between cells just before the final cut occurs during cytokinesis ^22^. Lipids have also been shown to be involved in several other aspects of cell division ^21^. The combination of known important roles, coupled with precise regulation of lipid identity and localization during division, suggested that cytokinesis would be a good model system to test the hypothesis that membrane-associated proteins in cells interact specifically with certain lipids. Applying lipid-trap mass spectrometry to membrane-associated proteins involved in cell division, we show that these proteins interact with specific lipids, and that these interactions change with the cell cycle. This work provides new insights into cytokinesis and lays the foundation for systematic analyses of interactions between membrane proteins and their associated lipids, and subsequent functional studies.

## Results

### Development of lipid-trap mass spectrometry - a technique to identify protein-lipid interactions

Our approach is to use cells expressing GFP-tagged proteins of interest, lyse the cells under detergent-free conditions to avoid dissociation of lipid-membrane bonds, immuno-precipitate the tagged protein, extract the lipids bound to the tagged proteins, and finally identify the lipids using lipidomic mass spectrometry (MS) (Fig. 1).

**Fig. 1:**
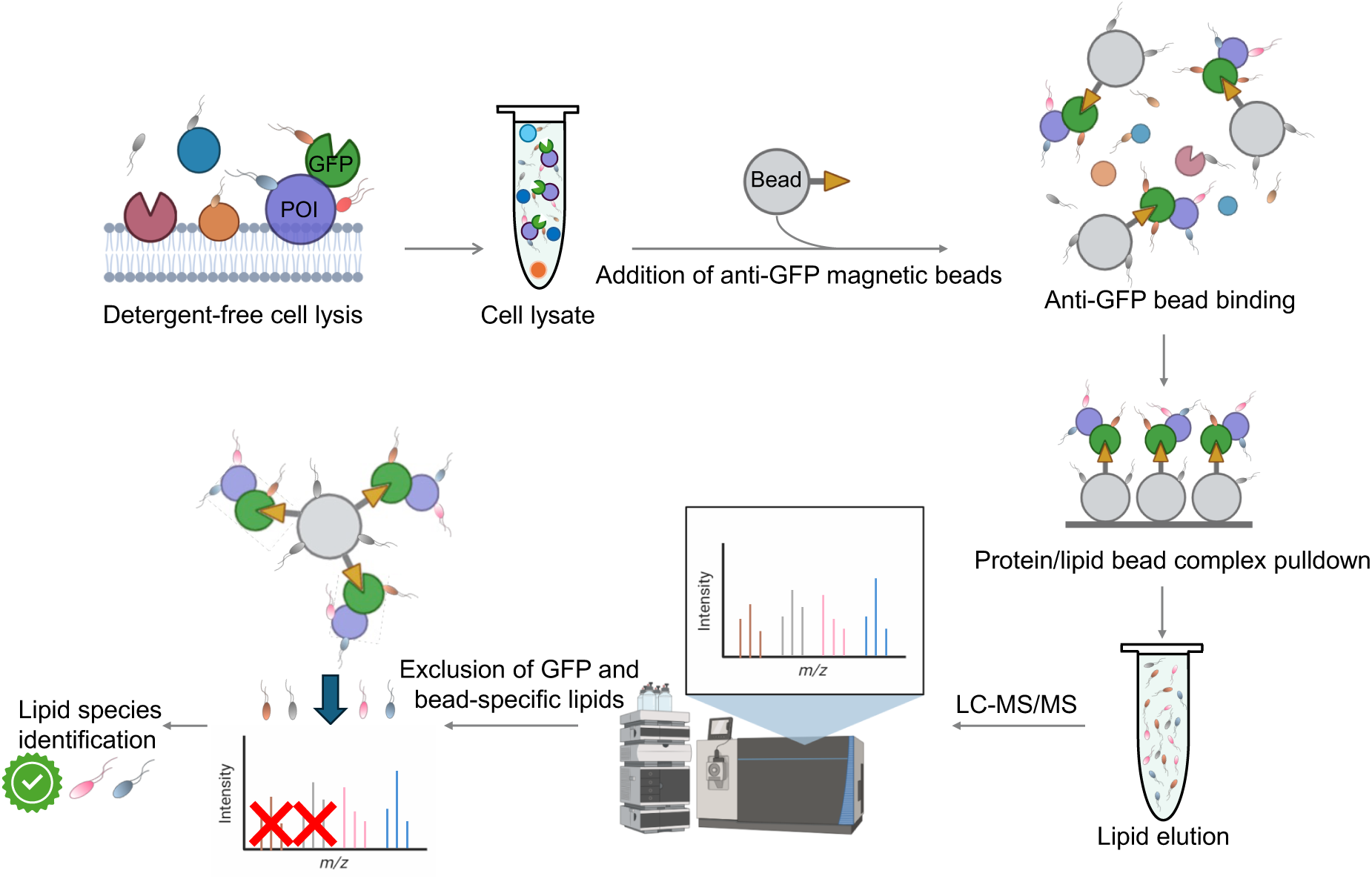
Experimental steps in lipid-trap mass spectrometry. A flowchart cartooning the identification of lipids bound to membrane-associated proteins of interest is shown.

The first step was to identify a protein tag that is reliable in pulldown experiments and is widely used, which is a pre-requisite for a generalizable and modular technique. We chose GFP – there are many well-characterized constructs available where GFP tags do not affect protein function. In addition, the correct localization of the protein of interest in the cell as well as attachment to the beads can be monitored by fluorescence microscopy. For our initial optimization we used a HeLa cell line expressing SEC61-GFP, a widely used marker of the ER (Extended Data Fig. 1a). GFP-Trap beads, which are magnetic agarose beads coated with a GFP antibody to pull down tagged proteins, are commercially available and commonly used ^23, 24^. We optimized lysis conditions by comparing sonication or mechanical disruption by vortexing with glass beads and passing the lysate through a syringe with a 27 G 1/2 needle (Extended Data Fig. 1b). A lysate obtained by sonication, followed by centrifugation at 3500 xg to clear large debris was added to GFP-trap beads, which resulted in fluorescent GFP signal distributed uniformly over the surface of the magnetic beads (Extended Data Fig. 1c and d). Using Nile red as a generic lipid label with high affinity for neutral lipids ^25^, we showed that GFP positive beads co-stained with the lipid dye, while there was little unbound membrane debris present (Extended Data Fig. 1c and d). After washing and isolation with a magnet, we extracted lipids from these beads with the organic solvent mixture CHCl_3_/MeOH (2:1, v/v) ^26^. Lipids were then separated by UPLC based on their size and hydrophobicity prior to being subjected to mass spectrometry. After the initial MS run, we performed statistical analysis to identify species (called features) that were enriched in LTMS pulldowns. We used membrane-localized myristoylated and palmitoylated GFP (MyrPalm-GFP) as a control for non-specific lipid binding (to GFP or beads) and excluded from subsequent analyses any lipids that were pulled down by controls alone (Fig. 1). Selected features with significant changes compared to control were then analyzed by tandem MS. LIPID MAPS ^27^, MS DIAL ^28^ and MS FINDER databases were used to match the lipid fragments from tandem MS and determine which lipid species corresponds to a selected feature.

### Lipid-trap MS identifies known lipid-protein interactions

Before applying LTMS to proteins involved in cell division, we validated this method using known lipid-protein interactions. First, we created a HeLa cell line stably expressing GFP tagged Lact-C2 (Fig. 2a). Lactadherin is a peripheral membrane protein involved in different functions, including stabilizing the phospholipid bilayer that surrounds triglyceride globules in breast milk and mediating phagocytosis of dying cells ^29^. Lactadherin has been shown to bind phosphatidylserine (PS) through its C2 domain (Lact-C2), which is used to visualize PS in cellular imaging studies ^30, 31^. We found that a PS species (PS 18:1_18:1) was indeed pulled down (Fig. 2b) (Supplementary Table 1). Several other lipids, mostly glycosylated ceramides, also associated with Lact-C2. Glycosylated ceramides are synthesized in the Golgi, as is a pool of PS, and then both are trafficked to the plasma membrane where glycosylated ceramides are usually found on the outer leaflet of the plasma membrane whereas PS is in the inner leaflet. These data provided the promising first hints that, as was our intention, we are isolating regions of lipids or areas of membrane associated with our protein of interest, rather than only non-annular interactors.

**Fig. 2:**
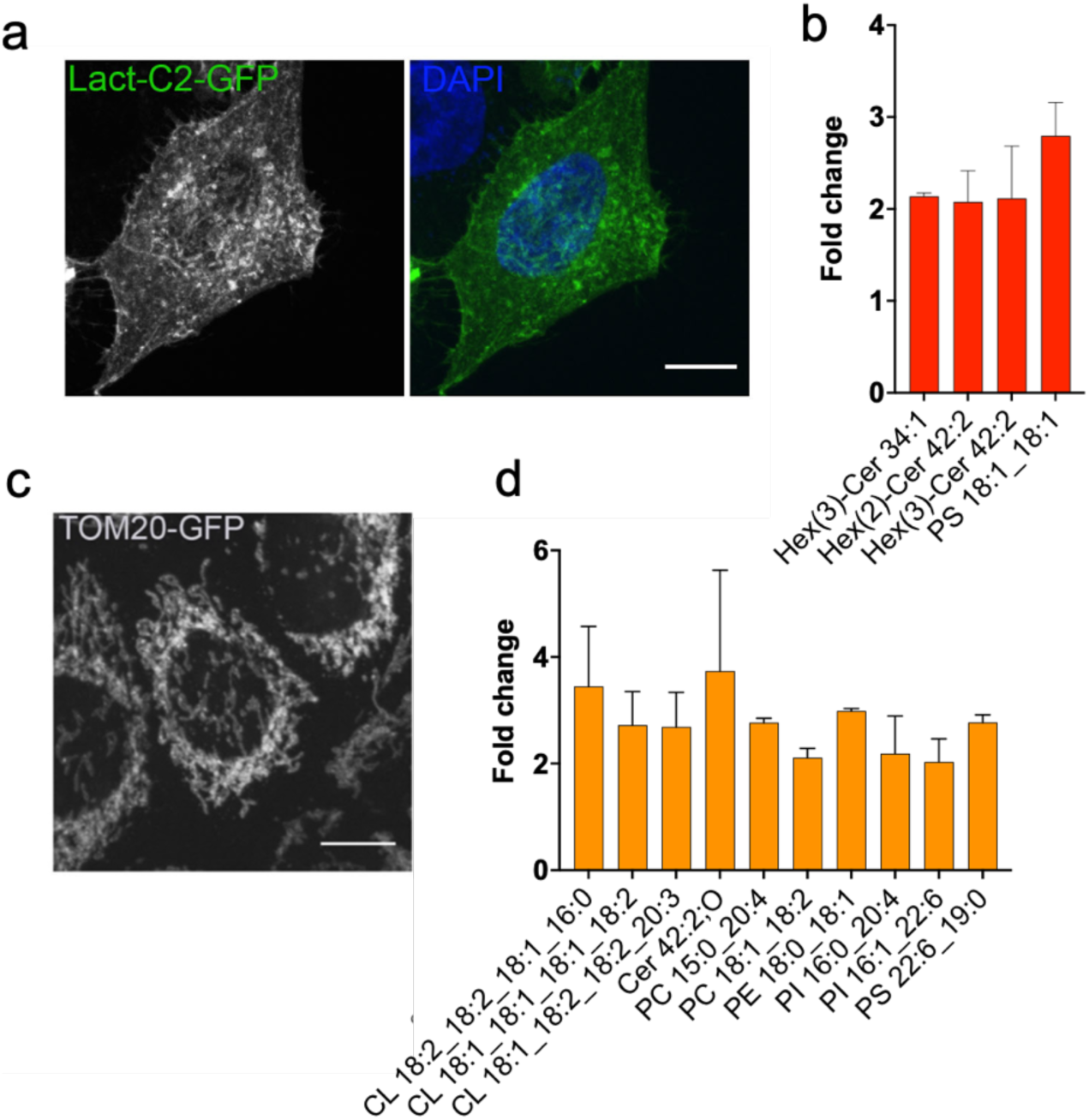
LTMS from GFP-Lact-C2 and TOM20-GFP. **a,** Representative confocal image of HeLa cell stably expressing GFP-LactC2 (green), fixed and stained with DAPI (blue) to visualize DNA, scale bar=10μm. **b,** Bar graph showing the fold increase of lipids detected with GFP-LactC2 LTMS compared to MyrPalm-GFP (control). Lipids with a fold increase ≥ 2 compared to MyrPalm-GFP are shown. Data represent mean value and standard deviation (SD), 3 replicates per experiment were run. N=3 experiments, p ≤ 0.05 for all lipids shown. **c,** Representative confocal image of HeLa cells stably expressing TOM20-GFP (grey), scale bar=10μm. **d,** Bar graph showing the fold increase of the lipids detected in TOM20-GFP LTMS compared to MyrPalm-GFP (control). Lipids with a fold increase ≥ 1.5 compared to the MyrPalm-GFP control are shown. Data represent mean value and standard deviation (SD), 3 replicates per experiment were run. N=3 experiments, p ≤ 0.05 for all lipids shown.

Next, we created a cell line stably expressing GFP-tagged TOM20. TOM20 is located in the outer membrane of mitochondria and is part of the mitochondrial protein translocation machinery (Fig. 2c). Applying LTMS, we identified several lipids associated with TOM20, including cardiolipins (Fig. 2d). Cardiolipins are unique to mitochondria, further showing that the lipids pulled down by our technique are specific to the protein of interest. Although cardiolipins are mostly found in the inner membrane of mitochondria, they are required for the function of the TOM complex, and specifically the interaction of TOM20 with other components of the complex ^32, 33^, confirming our finding. We also identified specific phospholipids, including PE, PI, PC and PS (Fig 2d, Supplementary Table 1), all of which are found in mitochondria ^34^. These data further support that we are identifying individual lipids as well as regions of membrane associated with our tagged proteins.

The lipids identified by LTMS from Lact-C2 and TOM20 were consistent for each protein across several repetitions, but different between the two proteins, minimizing the possibility that we are observing non-specific binding. Therefore, these validation experiments show conclusively that LTMS can successfully identify lipids associated with membrane proteins.

### The cytokinesis protein RACGAP1 binds to specific lipids during division

Having established that LTMS can identify lipids known to be associated with membrane proteins, we next investigated proteins involved in cytokinesis, starting with RACGAP1. RACGAP1 links the mitotic spindle to the plasma membrane and is essential for central spindle formation ^35^. It is localized at the intercellular bridge during cytokinesis and predominantly in the nucleus in interphase (Fig. 3a) ^36^. We chose to focus on this protein because it is essential for cytokinesis and has a known lipid interaction; a domain responsible for this interaction (C1) has been documented ^36^. To better understand any cytokinesis-specific interactions, we compared lipids bound to protein extracted from non-synchronized cells to those from cells synchronized at cytokinesis (~65% synchrony, Extended Data Fig. 2). As stable overexpression of RACGAP1 is deleterious to cells, we created stable cell lines expressing inducible full length RACGAP1-GFP ^36^ (Fig. 3a) as well as a mutant lacking the C1 lipid binding domain (Extended Data Fig. 3f). After expression of RACGAP1-GFP or RACGAP1-ι1C1-GFP was induced for 1 hour, we used LTMS to identify lipids associated with these proteins. It was previously shown that the C1 domain of RACGAP1 can interact with phosphatidylinositol phosphates (PIPs) PI(4,5)P_2_ and PI(4)P *in vitro* ^36^. As our standard lipidomic mass spectrometry protocol cannot reliably detect PIPs, we performed a PIP derivatization to investigate if RACGAP1 can bind to PIPs *in vivo.* This protocol cannot differentiate between PIP phosphorylation sites, e.g. PI(4,5)P_2_ vs. PI(3,4)P_2_ as they have the same molecular mass, but it can identify the side chain configurations within this lipid family, which previous studies were unable to do. We found that only full length RACGAP1 pulled down from cells synchronized at cytokinesis was able to bind PIPs. LTMS from non-synchronized cells or with constructs lacking the PIP-binding C1 domain showed no or small amounts of PIPs (Fig. 3b), confirming previous *in vitro* findings ^36^ and further validating our technique. From our control HeLa extract (prior to LTMS) we were able to identify 5 PIP and 4 PIP_2_ species with different side chain patterns, with PIP 18:0_20:4 and PIP_2_ 18:0_18:1 being the most abundant species (Supplementary Table 2). Interestingly, however, we could only detect a single PIP_2_ (PIP2 18:0_18:1, which is most abundant in HeLa extract), and a single PIP (PIP 18:0_20:2, which is least abundant in HeLa extract) species bound to RACGAP1 (Fig. 3b, Supplementary Table 1, Extended Data Fig. 4). Although there has been much research on the important signaling roles of PIPs ^20, 37^, there has been little emphasis on which specific PIP species might be involved. Our study shows that proteins in live cells distinguish between PIP species with high specificity and therefore presumably use specific PIP species for different purposes.

**Fig. 3:**
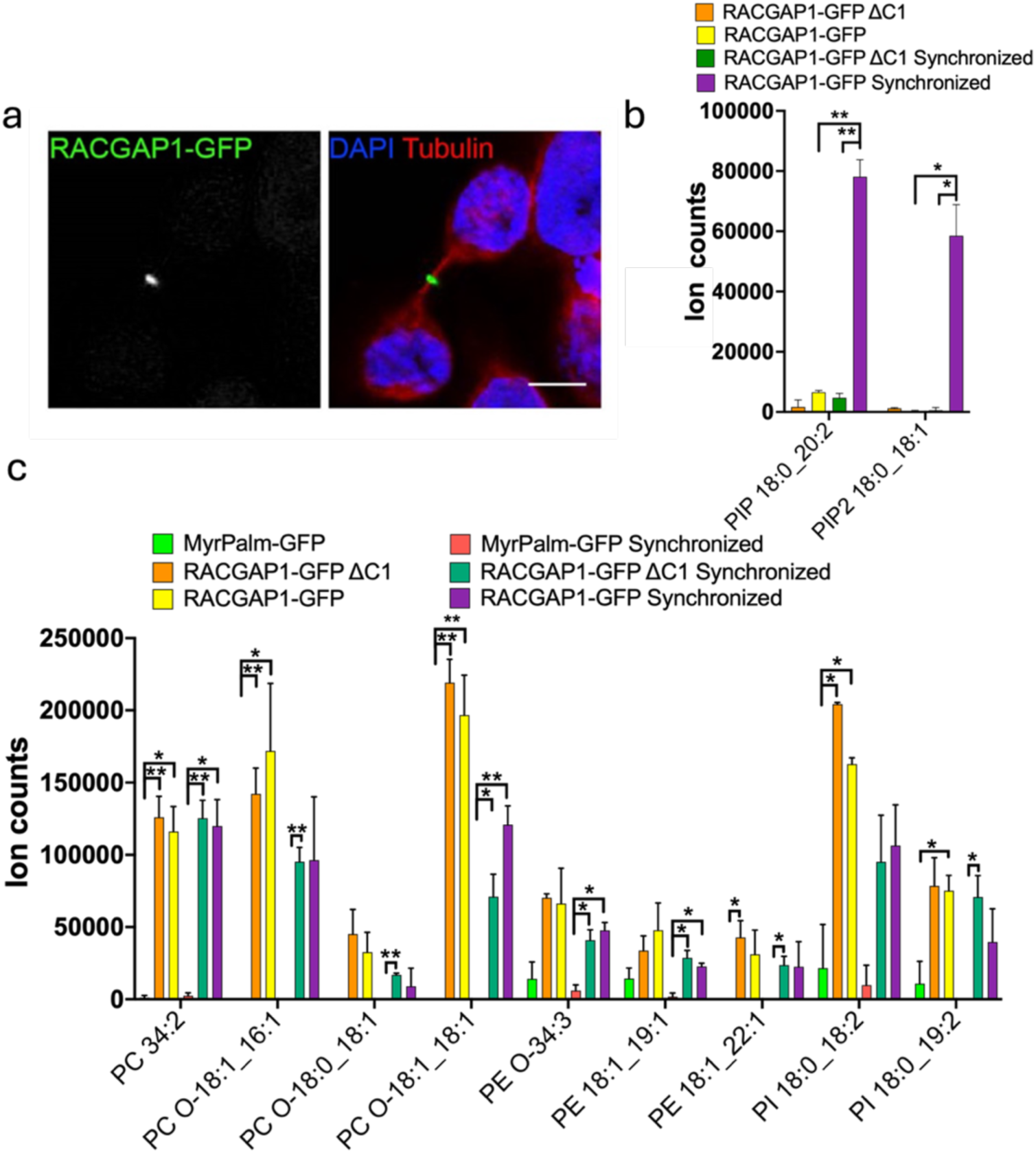
LTMS from RACGAP1-GFP. **a,** Representative confocal image of HeLa cell in cytokinesis stably expressing inducible RACGAP1-GFP (green), induced for 1h, fixed and stained with DAPI (blue) and anti-tubulin antibody (red) to visualize DNA and microtubules, scale bar=10μm. **b,** Bar graph showing the abundance of PIP 18:0_20:2 and PIP2 18:0_18:1 detected after LTMS of RACGAP1-GFP in wt and C1 deletion, in synchronized or unsynchronized cells. Data represent mean value and standard deviation (SD), 3 replicates per experiment were run. N=2 experiments. **c,** Lipid analysis from RACGAP1-GFP and RACGAP1-GFP ΔC1 (d) LTMS experiment in synchronized or unsynchronized samples. Statistical significance was assessed using multiple T-test based on false discovery rate (FDR) with the Two-stage linear step-up procedure of Benjamini, Krieger and Yekutieli, with Q = 5%. *, p ≤0.05, **, p ≤ 0.01; ***. Data represent mean value and standard deviation (SD), 3 replicates per experiment were run, N=2 experiments

In addition to analyzing PIPs, we also investigated if RACGAP1 and RAGAP1-ι1C1 bound other lipids (Fig. 3c, Supplementary Table 1). Several lipids were pulled down reproducibly, but independently of cell cycle state or inclusion of the C1 domain. These data suggest that RACGAP1 is able to interact with lipids specifically, but independently of the C1 domain. This may represent a general membrane attachment state that is then altered by specific interactions with PIPs via the C1 domain during cell division.

### Proteins of the ESCRT-III abscission machinery are associated with specific lipids

Next, we applied LTMS to two components of the ESCRT-III machinery required for the final cytokinetic membrane abscission: CHMP4B and CHMP2A. CHMP4B and CHMP2A are pivotal in cytokinesis as their depletion significantly perturbs abscission ^38, 39^. ESCRTIII components are dynamic and, in addition to an initial pool localized close to the midbody, a second pool, likely responsible for the final abscission, is found at the midbody periphery ^40^. The ESCRTIII complex is also essential for other biological functions such as endosomal sorting, viral budding and nuclear envelope sealing ^41, 42^. Similarly to CHMP4B (Fig. 4a), CHMP2A is localized to the midbody during cytokinesis (Fig. 4f), and, like CHMP4B, is recruited in all the main ESCRTIII-catalyzed processes ^43^. Despite acting at a similar timing during cytokinesis and both being necessary for abscission, CHMP4B and CHMP2A appear to have slightly different functions. CHMP4B seems to be the key component for the physical abscission of the intracellular bridge while CHMP2A is essential for CHMP4B localization and important for the recruitment of VPS4, the ATPase responsible for depolymerization of ESCRT-III ^44, 45^. We were interested to understand better how ESCRTIII interacts with membranes and if these two components that localize in the same membrane region interact with similar lipids.

**Fig. 4:**
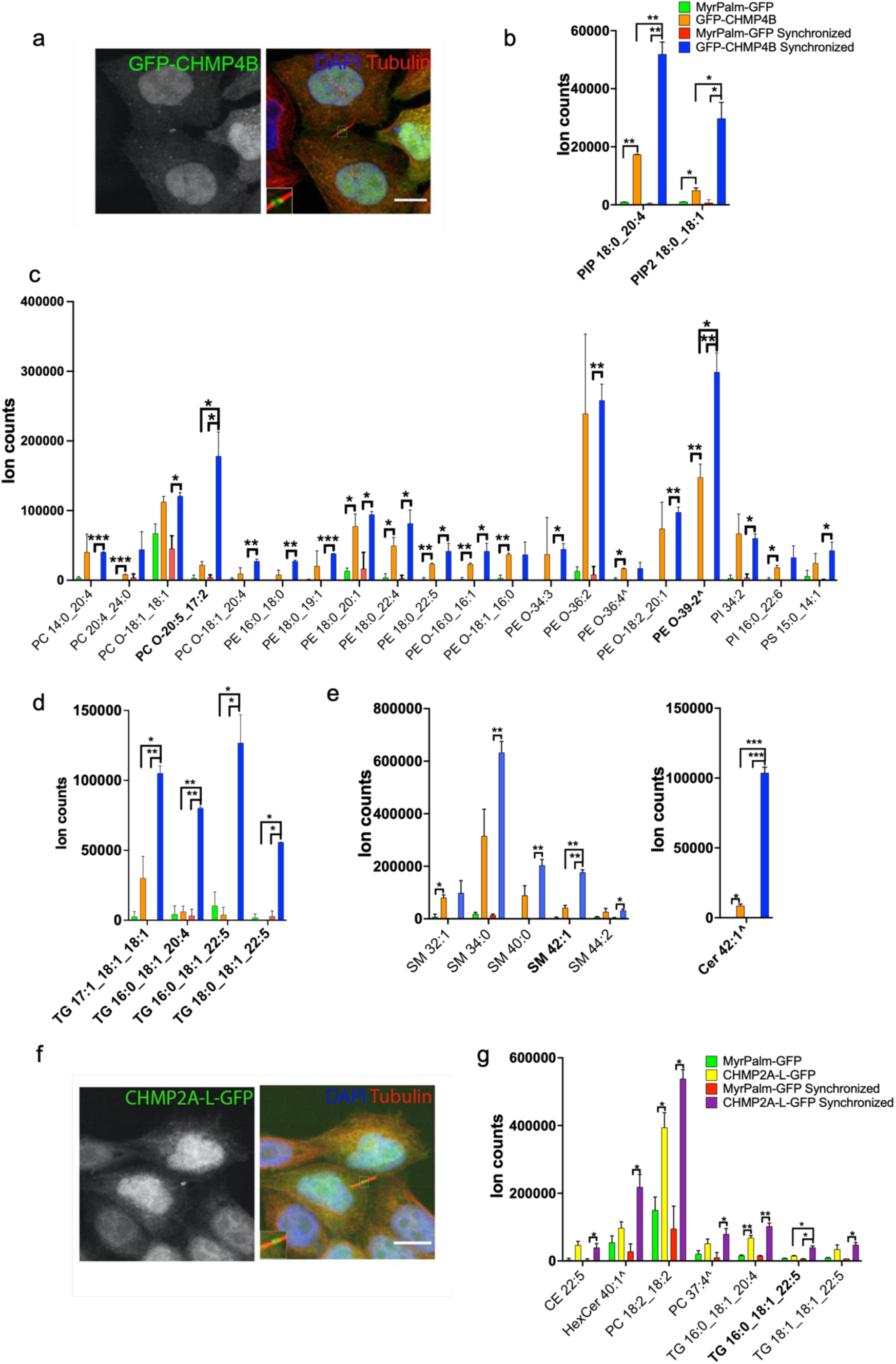
LTMS from GFP-CHMP4B and CHMP2A-L-GFP. **a,** Representative confocal images of HeLa cells stably expressing GFP-CHMP4B (green) fixed and stained with DAPI (blue) and anti-tubulin antibody (red) to visualize DNA and microtubules, scale bar=10μm **b,** Bar graph showing the abundance of PIP 18:0_20:4 and PIP_2_ 18:0_18:1 detected after pull-down of CHMP4B in synchronized and unsynchronized cells. **c-e,** Lipid analysis from GFP-CHMP4B LTMS in unsynchronized and synchronized samples showing phospholipids (**c**), triacylglycerols (**d**) and sphingolipids (**e**). Bold: increased in synchronized samples. **f,** Representative confocal images of HeLa cells stably expressing CHMP2A-L-GFP (green) in interphase and cytokinesis stained with DAPI (blue) and tubulin (red) to visualize DNA and microtubules, scale bar = 10μm. Inset: zoom with brightness enhancement of ICB. **g,** Lipid analysis from CHMP2A-L-GFP in synchronized and unsynchronized samples. Bold: increased in synchronized samples. Statistical significance was assessed using multiple T-test based on false discovery rate (FDR) with the Two-stage linear step-up procedure of Benjamini, Krieger and Yekutieli, with Q = 5%. *, p ≤ 0.05, **, p ≤ 0.01; ***, p ≤ 0.001, ****, p ≤ 0.0005. Data represent mean value and standard deviation (SD), 3 replicates per experiment were run. N=2 experiments. ^= tentative assignment with only retention time, m/z and adduct type due to low abundance.

*In vitro*, some experimental evidence suggests that CHMP4B has a higher affinity for membrane invaginations compared to planar membranes and that it binds preferentially to negatively charged membranes ^46^. In contrast, other findings show that CHMP4B and other ESCRT-III components do not have an affinity for negatively charged membranes on their own, but they may require ESCRT-I and ESCRT-II to coordinate their assembly ^47^. On the other hand, CHMP2A was shown to bind positively curved membranes and to be required for CHMP4B localization in membrane tube structures ^47^. These studies gave us important information about CHMP4B’s and CHMP2A’s preferred membrane conformation, but they were done using cell-free membrane models. Therefore, we decided to explore the nature of these interactions in cells.

The change in the localization of CHMP4B and CHMP2A between interphase and cytokinesis could suggest that the lipid species that these ESCRT-III components bind vary as cells progress through the cell cycle. For this reason, we synchronized cells at cytokinesis (Extended Data Fig. 2), as we did with HeLa cells expressing RACGAP1-GFP, and compared the LTMS data from dividing and non-dividing cells. Interestingly, we not only observed that specific lipids interact with CHMP4B and CHMP2A but also that some of these lipids were enriched in synchronized compared to unsynchronized samples. This shows that ESCRTIII alters its lipid binding according to its localization, suggesting lipids are involved in its regulation and/or function (Figs. 4b, 4c, 4d, 4e and 4g, Supplementary Table 1). These data also highlight the sensitivity of LTMS, and its utility in probing dynamic processes.

LTMS resulted in a variety of phospholipids, especially from GFP-CHMP4B (Figs. 4b and 4c, Supplementary Table 1), including some anionic phospholipids such as phosphatidylserines and phosphatidylinositols. This is consistent with the previous *in vitro* studies showing that CHMP4B has a higher affinity for negatively charged lipids ^46^. Neutral lipids species such as phosphatidylcholines, phosphatidylethanolamines and sphingomyelins, were also identified, suggesting that CHMP4B (and/or its protein complex partners) could also directly interact with or be surrounded by these lipid species. Ether phosphatidylethanolamines and ether phosphatidylcholines were also isolated (Fig. 4c, Supplementary Table 1). Ether phospholipids are characterized by an ether, rather than ester, bridge between their alkyl chain and their glycerol backbone and are major components of biological membranes (Appendix I) ^48^. However, apart from their structural roles and a potential role in membrane signaling and trafficking, it is not well known what functions they play in cells. Our result suggests that these lipid species interact with CHMP4B and potentially with other membrane-associated proteins. The most striking difference in LTMS of dividing and non-dividing cells was in triacylglycerols (TGs) and PIPs (Figs. 4b and 4d, Supplementary Table 1). Four different TG species were increased in the synchronized samples, suggesting a possible specific interaction of CHMP4B with these lipids during cytokinesis. In addition, after performing the derivatization protocol, we detected PIP 18:0_20:4 and PIP_2_ 18:0_18:1. PIP2 18:0_18:1 was the same species identified from RACGAP1 suggesting that this PIP_2_, the most abundant detected in the total HeLa extract (Supplementary Table 2), might have a role in the localization of membrane-associated cytokinetic proteins at the midbody. This result confirmed previous *in vitro* studies showing that snf7, the yeast orthologue of CHMP4B, binds PIPs ^49^. Another important difference between synchronized and unsynchronized samples was in Cer 42:1, a lipid with a characteristic sphingosine backbone. Interestingly, another sphingolipid, SM 42:1, was increased in synchronized samples (Fig. 4e). Sphingomyelins are ceramides linked through a phosphodiester bond to a choline head group. SM 42:1 has the same number of carbons and double bonds as Cer42:1, suggesting it might be a related species, although our mass spectrometer is not capable of proving this conclusively as it cannot determine the location of double bonds. These data might indicate that CHMP4B, when isolated from dividing cells, has a higher affinity for sphingolipids with the specific 42:1 configuration. Along with TGs, SM 42:1 and Cer 42:1, PC O-20:5_17:2 and PE O-39:2 were also observed to be increased in synchronized samples. The open question now is how this variety of lipids contributes to CHMP4B function during cytokinesis.

LTMS identified fewer lipids from CHMP2A: two PC species, especially PC 18:2_18:2, a cholesterol ester species, CE 22:5, and a glycosylated ceramide, HexCer 40:1, were enriched. This indicates that CHMP2A directly interacts with these lipid species or that it is localized in a subregion rich in these lipids (Fig. 4g, Supplementary Table 1), suggesting that this construct of CHMP2A might localize in a different membrane environment compared to CHMP4B. Interestingly, unlike CHMP4B, CHMP2A interacted exclusively with neutral lipids, and did not interact with PIPs. This supports the idea that CHMP2A does not have an affinity for negatively charged lipids such as anionic phospholipids ^47^. One caveat is that CHMP2A fused to a flexible linker followed by GFP, CHMP2A-L-GFP, was used for the experiment. The presence of this linker is required to observe the expected localization of CHMP2A at the midbody ^50^, but may have altered its molecular interactions. The most interesting result for CHMP2A was in the TGs it associated with. As for CHMP4B, there was a significant difference in abundance between synchronized and unsynchronized samples for a specific TG species, TG 16:0_18:1_22:5. We detect 92 ester and ether TGs in our control Hela lipidome (Supplementary Table 2), with TG 16:0_18:1_22:5 making up ~ 2.7% of all TGs. However, more abundant species (which collectively make up ~60% of TGs) were not bound to CHMP4B or CHMP2A, suggesting these proteins can discern between TG species. The fact that two independent experiments with different complex partners pull down the same lipid suggests that TGs might be involved in the interaction of ESCRT-III with membranes during cytokinesis, although it is unclear where in the cell they would meet as TGs are usually found in lipid droplets or the endoplasmic reticulum.

Lipid droplets are organelles that store neutral lipids including TGs. To investigate if CHMP4B interacts with TGs in lipid droplets, we transiently transfected HeLa cells stably expressing GFP-CHMP4B with BFP-Livedrop ^51^. Livedrop is a hydrophobic hairpin sequence derived from GPAT4. GPAT4 accumulates on lipid droplets as they form in the endoplasmic reticulum ^51^ and is a marker for nascent lipid droplets. However, we did not observe a strong colocalization between CHMP4B and Livedrop, suggesting that CHMP4B interacts with TGs in other membrane structures or the cytosol (Fig. 5a). To investigate this, we used ultracentrifugation to separate different cell components (Fig. 5b) and performed LTMS. CHMP4B was equally distributed in the fractions and pellet (Extended Data Fig. 5), yet bound to TGs primarily in the pellet fraction from dividing cells (Figs. 5b and c). This suggests that CHMP4B interacts with TGs in membrane structures in dividing cells, supporting a role during cell division. These data again highlight the utility and versatility of LTMS, permitting analysis of sub-cellular fractions from live cells.

**Fig. 5:**
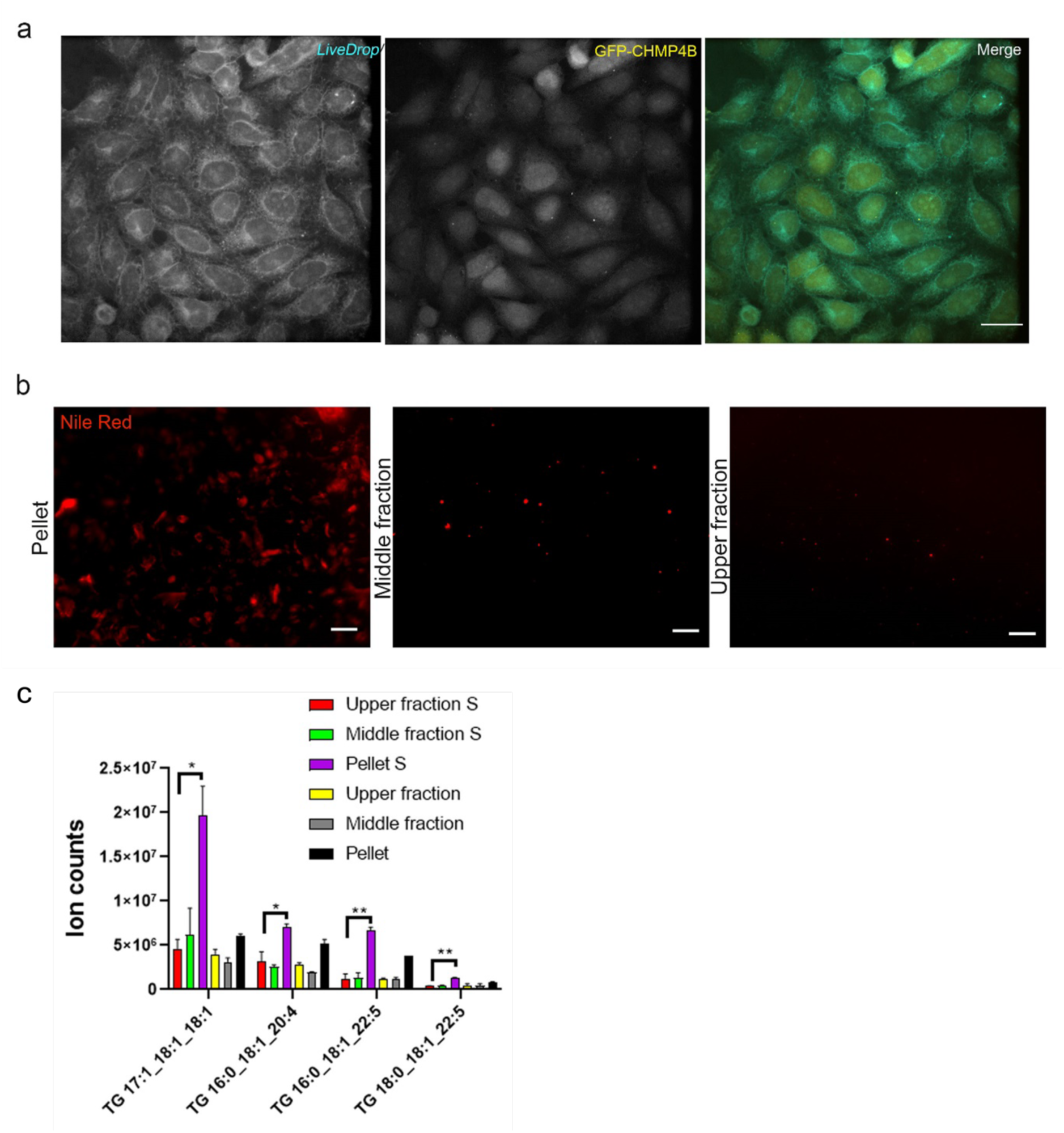
CHMP4B interacts with TGs in membranes. **a,** Representative confocal images of HeLa cells stably expressing GFP-CHMP4B (yellow) transiently transfected with BFP-*LiveDrop* (cyan) ^52^, scale bar=20μm. **b,** Microscopy images of membrane fragments stained with Nile Red after ultracentrifugation, scale bar=5μm. **c,** Bar graph showing the abundance of the TGs species binding CHMP4B after ultracentrifugation in pellet, middle and upper fractions, in synchronized (S) and unsynchronized samples. Statistical significance was assessed using multiple T-test based on false discovery rate (FDR) with the Two-stage linear step-up procedure of Benjamini, Krieger and Yekutieli, with Q = 5%. *, p ≤ 0.05, **, p ≤ 0.01. Data represent mean value and standard deviation (SD), 2 replicates were run.

## Discussion

We introduce here a technique, named lipid-trap mass spectrometry (LTMS), that allows systematic analysis of protein-lipid interactions from dynamic mammalian cells and subcellular fractions. This versatile technique can be applied to any integral or peripheral membrane protein, including those residing in organelles, as long as it can be tagged with GFP or a similar tag for which affinity reagents are available. Developing a new method to investigate protein-lipid interactions is crucial as lipids and membranes are essential in numerous biological processes. Here, we present a model study where we investigate protein-lipid interactions during cell division.

We show that membrane-associated cytokinetic proteins bind specific lipids, contributing to our understanding of why cells regulate their lipidomes during division with high precision ^22, 53^. Key cytokinesis proteins RACGAP1 and CHMP4B both bound distinct lipids in dividing compared to non-dividing cells, further supporting the observation that membranes are highly dynamic, yet specifically staged, during cell division. Gratifyingly, LTMS identified PIPs as interactors, as would be expected from the literature, although with unprecedented chemical precision as the side chain configurations of key PIPs were not previously known. Our work also found several unexpected lipid-protein interactions with a range of different lipid species. This includes specific phospholipids and sphingolipids and, surprisingly, a small number of TGs. These TGs are not the most abundant species found in HeLa, suggesting that different TGs have distinct binding partners and therefore cellular roles. There is much research to be done to understand these interactions, as the field currently does not have a strong understanding of how cells regulate and mediate the coordinated synthesis, metabolism, transport and interactions of the large number of lipids it produces. LTMS will enable investigations towards this goal.

An advantage of LTMS is that it allows the identification of regions of lipids surrounding a protein of interest. This is important because annular and bulk lipids likely influence protein function by creating a very local membrane environment with specific physical and signaling properties, which may be needed for function, including recruitment of proteins and lipids, specific protein conformations or interactions. This local environment can also include other proteins, which could be identified by blotting or proteomics if needed for functional analysis. We expect that LTMS can be expanded to distinguish between annular, non-annular and bulk lipids by adding low concentrations of detergent and/or changing the pH or ionic strength of the lysis and washing buffers, concentrating lipids that bind more tightly to the protein of interest ^54^. While we present cellular studies in this initial report, we anticipate that LTMS will have applications in biophysical and structural investigations of membrane proteins. Currently, proteins in cell-free studies are often embedded in model membranes that include a handful of standard lipid species like DOPC or POPC. Applying our lipid isolation protocol would allow reconstitution of a membrane that resembles the protein’s native cellular environment more closely.

This initial report of LTMS identified lipids that surround and interact with GFP-tagged proteins. However, the simple and modular nature of LTMS makes it versatile, and it should be applicable to proteins tagged with other widely used tags, such as FLAG, HA, GST, Halo or SNAP tags, which are frequently used in cell biology ^55, 56^. It may also be possible to adapt immunoprecipitations if the specific system allows a largely detergent free workflow. We therefore hope, and expect, that LTMS will become a new tool in our arsenal of understanding the lipids and proteins of many different membranes. A new door has been opened for exploring the role of lipids in cell biology.

## Materials and Methods

### Cell culture

HeLa cells, a gift from Prof. Jeremy Carlton, were authenticated by STR profiling (Eurofins MWG). HEK293GP and HEK293T cells were from Clontech (Takara Bio). All cell lines were regularly confirmed mycoplasma-free by PCR assay (EZ-PCR^TM^ Mycoplasma Test Kit, Biological Industries) and DAPI staining. All cell lines were cultured under standard conditions at 37**°**C, 5% CO_2_ and were maintained in DMEM medium (Gibco, Thermo Fisher Scientific) with addition of 10% Fetal Bovine Serum (FBS) (Gibco, Thermo Fisher Scientific) and 100 U /100 µg Penicillin/Streptomycin/L (P/S) (Sigma Aldrich) (complete growth medium). Medium of stable cell lines was supplemented with Puromycin 0.5 μg/mL or Geneticin (G418) 400 μg/mL (Gibco, Thermo Fisher Scientific). HeLa expressing inducible RACGAP1-GFP or RACGAP1-GFP ΔC1 were further supplemented with Doxycycline 1 μg/mL (Sigma Aldrich) for experiments.

### Cell cycle synchronization

HeLa cells stably expressing the GFP-tagged protein of interest were counted and plated at a density of 4×10^6^ cells per T75 flask. 3 replicates were used for each experiment. The day after, 100 ng/mL nocodazole (Sigma Aldrich) was added for 12 h. Medium was removed, and mitotic cells were collected by mitotic shake-off. Cells were then transferred to Falcon tubes and centrifuged at 200 x*g*. Pellets were washed with Dulbecco’s PBS to remove the residual nocodazole and resuspended in medium. Cells were then re-plated on 10 cm dishes and released from mitotic arrest for 120 min before the LTMS pull down. HeLa expressing inducible RACGAP1-GFP and RACGAP1-GFP ΔC1 were supplemented with Doxycycline 1 μg/mL (Sigma Aldrich) during the release,

### Immunofluorescence

Cells were plated on 11 mm glass coverslips and fixed using 4% paraformaldehyde (PFA) (Alfa Aesar) in PBS. Cells were fixed for 20 min, permeabilized using 0.1% Triton X-100 in PBS for 5 min and washed twice with PBS/0.5% BSA/20 mM glycine/0.1% NaN_3_ to neutralize unreacted aldehydes for 15 min. Samples were then stained with primary antibodies overnight at 4°C and secondary antibodies for 2h at room temperature in blocking buffer (PBS/1% BSA (Apollo Scientific) /0.1% NaN_3_ (Sigma Aldrich)). Confocal images were acquired at the Nikon Imaging Centre, King’s College London. Acquisition was performed on an inverted Nikon Eclipse Ti microscope equipped with a Yokogawa CSU-X1 spinning disk unit and a Andor Neo sCMOS camera. The microscope was equipped with a 405 nm laser for DAPI excitation, a 488 nm laser for FITC excitation and a 561 nm laser for TRITC excitation. A 40X air lens was used for acquisition. Membrane fragments after ultra-centrifugation were imaged using an inverted Nikon Eclipse microscope using widefield epifluorescence (Ti-E) equipped with a Cool SNAP HQ 2, DS-Fi2 Color CCD camera using a 20x objective.

### Western blot

Laemmli lysis-buffer 1.5X without bromophenol blue or DTT (3% SDS Sodium Dodecyl Sulphate (Sigma Aldrich), 15% glycerol, 0.094M Tris-HCl pH 6.8 (Fisher Scientific)) was used to lyse cells. Plates were incubated on a heating block for 10 min at 100°C. After removing the plates from the heating block, each sample was passed through a 25G needle (Terumo) 10 times. 3x Sample Buffer (6% SDS, 30% glycerol, 0.003% bromophenol blue, 0.2M Tris/pH 6.8) and 0.3M DTT were added to cells to give a final 1x concentration and then lysates were transferred to microcentrifuge tubes and heated for 5 min at 100°C. Samples were stored at −20°C until further analysis. Proteins were resolved using SDS–PAGE and transferred onto a 0.22 μm nitrocellulose membrane. Proteins were then incubated with antibodies of interest in 5% milk overnight at 4°C. The following day, nitrocellulose membranes were incubated with secondary HRP-conjugated antibodies (Jackson ImmunoResearch) and imaged.

### Antibodies and fluorescent dyes

The following primary antibodies were used: mouse anti-αTubulin (Thermo Fisher 236 10501, immunofluorescence microscopy 1:1000), mouse anti-RACGAP1 (Everest Biotech EB05315, western blot 1:1000), mouse anti-GAPDH (Proteintech10494-1-AP anti-GFP, western blot 1:5000), rabbit anti-GFP (Abcam ab6556), Rabbit and mouse horseradish peroxidase (HRP) conjugated secondary antibodies (Stratech-Jackson Immunoresearch), Alexa 488 rabbit and mouse (Jackson ImmunoResearch AB_2340846 and Jackson ImmunoResearch AB_2338871), Alexa 594 rabbit and mouse (Jackson ImmunoResearchAB_2313584 and Jackson ImmunoResearch AB_2338046). DNA was stained with 1 mg ml^−1^ 4′,6-diamidino-2-phenylindole (DAPI; 1:1000) and neutral lipids with 9-(Diethylamino)-5H-benzo[a]phenoxazin-5-one (Nile Red, Thermo Fisher N1142, 1:100).

### Generation of stable cell lines

Lentiviral and retroviral particles were generated using HEK293T or HEK293GP cells, respectively. Lentiviral transfection mixtures were prepared by adding 1700 ng plasmid of interest, 1700 ng pVSVG (Addgene, #138479), 1700 ng psPAX2 (Addgene, #12260) and 4μg/ml polyethyleneimine (Sigma-Aldrich) in a total volume of 200 μL DPBS. Retroviral transfection mixtures were prepared by adding 1200 ng/ml plasmid of interest, 800 ng/ml pVSVG and 1μg/ml polyethyleneimine in a total volume of 200 μL DPBS. The plasmids were then transfected into HEK293T or HEK293GP cells. HEK293GP cells constitutively express gag and pol proteins for retrovirus packaging and psPAX2 was not added to the transfection mixtures. After 4 h, the medium was replaced, and cells were left for 48 h to produce the viral particles. Medium was harvested and viral particles were then transduced in HeLa cells.

### Plasmid constructs

pCW57-GFP-2A-MCS was a gift from Adam Karpf (Addgene plasmid # 71783), Lact-C2-GFP was a gift from Sergio Grinstein (Addgene plasmid # 22852), mCherry-TOMM20-N-10 was a gift from Michael Davidson (Addgene plasmid # 55146). RACGAP1 cDNA was a gift from Prof. Francis Barr ^57^. pNG72-CHMP2A-LAP-GFP was a gift from Prof. Juan Martin Serrano ^50^. pAG138-BFP-Livedrop and pCMS28-GFP-CHMP4B were a gift from Prof. Jeremy Carlton. pcW57 vector was a tetracycline/doxycycline-inducible lentiviral vector, Tet ON. pcW57-RACGAP1-GFP was created by digesting pcW57 with NheI and BamHI and ligating the GFP insert. The pcW57-GFP vector was then further digested with NheI and EcoRI to insert RACGAP1. RACGAP1 cDNA was amplified by PCR using primers 5’ AAAAAAGCTAGCACCATGGATACTATGATGCTGAATGTG 3’ and 5’ AAAAAAGAATTCCTTGAGCATTGGAGAAGC 3’. A Q5 site-directed mutagenesis kit (NEB) was used to delete RACGAP1 C1 domain. The following primers were used for deletion: 5’ CCTACCCTGATAGGAACAC 3’ and 5’ GCGCATCCCTCCATTACT 3’. Ligation was performed with a T4-DNA-ligase kit (M0202, NEB). The constructs were verified by DNA sequencing.

### LTMS pulldowns

Non-synchronized Hela cells stably expressing the GFP-tagged protein of interest were counted and plated at a density of 10^6^ cells per 10cm dish. 3 replicates were used for each experiment. The following day cells were transferred to ice. Synchronized cells were released from mitotic arrest for 120 minutes before transfer to ice. The same protocol was then applied for both synchronized and non-synchronized cells. Media was removed, and dishes were washed with DPBS. Cells were then harvested in PIPES (Sigma) buffer 20mM pH 6.8 (+ 1xProt inhibitor) (Roche) by scraping and transferred to pre-cooled Eppendorf tubes on ice. Cells were lysed through probe sonication (Sonics) by alternating 20 seconds on-cycle and 30 seconds off-cycle on ice for a total of 3 min at 30KHz. Samples were then centrifuged at 3500 x*g* for 10 min at 4°C. An aliquot of each cell lysate was saved as input lane for western blotting. GFP magnetic agarose beads (Chromotek) were washed twice with PIPES buffer before equilibration. Cell lysate was transferred to new Eppendorf tubes with 40 μL of GFP magnetic beads and equilibrated for 1 h at 4°C in tube rotators. After equilibration, Eppendorf tubes were placed in a magnetic rack, supernatants were discarded and beads were washed twice with PIPES buffer. Beads were subjected to lipid extraction for analysis of bound lipids or, alternatively, beads were resuspended in 3x sample buffer with DDT and boiled for 10 min at 95°C to dissociate proteins for Western blot analysis.

### Ultracentrifugation

To isolate cytoplasm and membrane fragments, samples were centrifuged in Eppendorf tubes at 720 xg for 5 min. Supernatants were transferred in new Eppendorf tubes on ice and centrifuged at 10,000 xg for 5 min. The new supernatants were transferred in ultracentrifuge Beckman tubes (Gibco, Thermo Fisher Scientific). Samples were then centrifuged at 100,000 xg for 1h with a Beckman Coulter Optima MAX XP Ultracentrifuge. Pellets were resuspended in PIPES. The middle and upper fractions were processed separately. The different fractions were then transferred to new Eppendorf tubes with 40 μL of GFP magnetic beads and the steps in the section above were followed.

### Lipid extraction

After washing the magnetic beads, samples were transferred to glass tubes. Aqueous phases were removed by using the magnetic rack flipped vertically.

For neutral lipid extraction, lipids were extracted using the method developed by Folch *et al.* ^26^. 250 μL of CHCl_3_/MeOH (2:1, v/v) was added to the magnetic beads. Glass tubes were vortexed twice for 60 seconds and the CHCl_3_/MeOH mixture was transferred to chloroform-resistant Eppendorf tubes.

For extraction of phosphoinositides, a modified version of the Folch method was used following the protocol by Mucksch et al. ^58^. 726 μL of CHCl_3_/MeOH/ HCl (1 M) (40:80:1, v/v/v) was added to the magnetic beads and the glass tubes were vortexed for 15 min. 720 μL of CHCl_3_ was added and the tubes were vortexed for 5 min. 354 μL of HCl (1 M) was then added and beads were vortexed for 2 min. The tubes were centrifuged at 1000 x*g* for 5 min and the lower phase, containing the phosphoinositides, was transferred to fresh Eppendorf tubes. 726 μL of CHCl_3_/MeOH/ HCl (1 M) (40:80:1, v/v/v) was added, and samples were vortexed for 10 seconds. Samples were centrifuged at 1000 x*g* for 5 min and the lower phase was transferred to the Eppendorf tubes from the first extraction.

Organic phases were evaporated on a heat block at 37°C under a constant N_2_ flow. After resuspension in the loading buffer, samples were analysed by LC-MS.

To exclude any unspecific binding to the beads or unspecific lipid association, we used HeLa cells stably expressing MyrPalm-GFP as a control. Samples were normalized by the number of cells plated.

### Phosphoinositide derivatization

For phosphoinositide derivatization, synchronized and non-synchronized cells were counted and plated at the same cell density as used for the Folch extraction. A reaction was performed following the protocol by Hille *et al.* ^59^. 90 μL of MeOH/CH_2_Cl_2_ (4/5, v/v) was added to resuspend the dried samples. 10 μL of TMS-diazomethane in hexane (2 M) (Sigma Aldrich) was added to each extract. The reaction was left to proceed for 30 min at room temperature. 20 μL of glacial acetic acid was then added to quench the excess of TMS-diazomethane. A beaker with glacial acetic acid was placed in the hood to prevent inhalation of possible volatile TMS-diazomethane. This reaction releases N_2_ (gas) and can be vigorous if a high excess of TMS-diazomethane is neutralized. Organic phases were dried and resuspended in loading buffer. Samples were analysed by LC-MS.

30 pmoles of PI(4)P and 60pmoles of PI(3,4)P_2_ brain lipid analytical internal standards (ISDs) were used in runs to compare exact mass, retention time and DAG fragments. ISDs were purchased from Avanti Polar Lipids (LIPID MAPS MS Standards; Avanti Polar Lipids, Alabaster, AL, USA). ISDs were dissolved in CHCl_3_/CH_3_OH/H_2_O (20:9:1, v/v/v) and further diluted to give a concentration of 2 ng/μL.

### Lipidomics analysis by LC-QTOF

Samples were resuspended in 100 μL loading buffer (1:1:2 water:acetonitrile (LC-MS grade):isopropanol (LC-MS grade)) (Honeywell, Fisher Scientific and Merck) and centrifuged at 6500 x*g* for 2 min to pellet any particulates to avoid column clogging due to injection of beads or precipitated proteins. 90 μL of each sample were then transferred to autosampler vials (Agilent).

An Agilent 1290 Infinity II UHPLC system coupled to an Agilent 6550 iFunnel Q-TOF mass spectrometer (Agilent Technologies) was used for the reversed phase-ultra-high performance liquid chromatography-mass spectrometry analysis (RP-UHPLC-MS). The UHPLC system was equipped with an Acquity UHPLC CSH C18 column (100 x 2.1mm, 1.7μm) (Waters). The column was maintained at 65°C and at a flow rate of 0.6 mL/min. The mobile phase was characterized by a mixture of (A) 60:40 (v/v) acetonitrile /H_2_O and (B) 10:90 (v/v) acetonitrile/isopropanol. For analysis in positive mode, the mobile phase was supplemented with 10mM ammonium formate and 0.1% formic acid, whereas for analysis in negative mode 10mM ammonium acetate was added to the mobile phase. Samples were run in a gradient of mobile phase A and B. The UHPLC-MS method was optimized with minor modifications from Cajka and Fiehn ^60^. Between injections, the needle was washed with 100% isopropanol.

UHPLC analytical gradient was set to: B from %15 to %30 (0-2 min): B from 30% to 48 % (2-2.5 min); B from 48% to 82% (2.5-11 min); B from 82% to 99% (11-11.5 min) and B kept on 99% for 4 minutes. In 0.5 mins B returned to its initial conditions and column was equilibrated for 3 minutes for the next run.

The QTOF was calibrated in the mass range 50-1700 *m/z* and spectra were acquired in centroid mode. After MS1 analysis, the samples were aligned and statistically analyzed as described in the last paragraphs. An inclusion list with *m/z* values and retention times of the features of interest was then created. The selected features were further investigated by targeted MS2. The collision energy was set at 25 eV and −20 eV for positive and negative mode, respectively. Agilent tune mix (mass resolving power ~10,000 FWHM) was used to tune the instrument. The reference solution to correct small mass drifts was composed of m/z 121.0509, *m/z* 922.0007 in positive mode, and *m/z* 119.0360, *m/z* 966.0007 (formate adducts), *m/z* 980.0164 (acetate adduct) in negative mode.

### Instrument Parameters

The following parameters were used for both positive and negative mode analyses: SheathGasFlow 11, SheathGasTemp 350, Nebulizer (psig) 35, Gas Flow (l/min) 14, and Gas Temp (°C) 200.

### Scan Source Parameters

The following parameters were used for both positive and negative mode analyses: OctopoleRFPeak 750, Skimmer1 0, Fragmentor 350, Nozzle Voltage (V) 1000 and VCap 3500.

### Lipidomics data analysis

MassHunter Profinder Software (version B.08.00 and B.10.00, Agilent Technologies) was used to process and analyze the data. For feature extraction, the “Batch Recursive Feature Extraction for Small Molecule” option was chosen. Between the possible adducts, H^+^, Na^+,^ and NH_4_^+^ were selected for positive mode, and H^−^, CH_3_COO^−,^ and HCOO^−^ were selected for negative mode. For the correct peak alignments, retention time (RT) span was set to 0.500 min and mass tolerance was set to 20 ppm. A retention time window between 0.500 and 12.000 min was analysed. Features originating from the control (beads) and solvents were removed.

Processed data were exported as CEF files from MassHunter Profinder and were imported to Mass Profiler Professional (MPP, Agilent Technologies) for statistical analysis. Replicates for each experimental condition were grouped. Features that were not present in at least 80% of samples were excluded and the remaining features were subjected to statistical analysis by ANOVA (p<0.05) with post hoc Tukey HSD test for each group compared to MyrPalm-GFP (control). According to the MPP analysis, raw data were checked again with MassHunter Profinder. Then, the (*.csv) files containing *m/z*, retention time, and area of the features were exported from MassHunter Profinder and the areas were further analysed with GraphPad Prism using T-tests based on false discovery rate (FDR) with the Two-stage linear step-up procedure of Benjamini, Krieger and Yekutieli, with Q = 5%.

MassHunter Qualitative Analysis Software (version B.07.00, Agilent Technologies) was used to generate extracted ion chromatograms (EIC) for lipid species with known *m/z* and retention times, such as derivatized phosphoinositides. The areas of the EICs were then exported and analysed with GraphPad Prism using T-tests based on false discovery rate (FDR).

Selected features with significant changes, compared to the control, were analysed by MS2. LIPID MAPS ^27^, MS DIAL ^28^, and MS FINDER databases were used to match the lipid fragments. If the interpretation of the MS2 spectrum was difficult due to background noise, total HeLa extracts were used to target the features. In total HeLa extracts, features are more abundant, which allows reduction of the background noise and assignment of the lipids with more confidence. Otherwise, accurate mass (<5 ppm deviation from expected mass), retention time, and the pattern of adduct formation were the parameters considered for the assignment.

Lipid annotations were performed following the guidelines of the Lipidomics Standard Initiative (LSI) (https://lipidomics-standards-initiative.org/).

### Lipidomics statistics

Mass Hunter Profinder Software (version B 10.00) was used for data processing. The ‘Batch Recursive Feature Extraction for Small Molecule’ option was used to select the features of interest. H^+^, Na^+^ and NH_4_^+^ adducts were selected for positive mode and H^−^, CH_3_COO^−,^ and HCOO^−^ adducts were selected for negative mode. Retention time (Rt) span was set to 0.500 min and mass tolerance to 20ppm.

CEF files were exported from Mass Hunter Profinder Software and imported in Mass Profinder Professional (MPP) for the statistical analysis. Features were statistically analysed by ANOVA (p<0.05) with posthoc Tukey HSD test for each group compared to MyrPalm-GFP (control). After deleting the non-significant features in Mass Hunter Profinder Software, the *(csv) files containing m/z, retention time, and intensity of features were exported. The intensity was analyzed with GraphPad Prism using T-test based on false discovery rate (FDR).

The features were further analyzed by MS2. LIPID MAPS, MS DIAL, and MS FINDER databases were used for the in-silico analysis of the fragments.

## Data Availability

All data supporting this study are available within the paper and its Supplementary Information. This includes MS/MS data used to identify the lipid species reported in the manuscript (Appendix I). Should any further raw data be required in a different format, they are available from the corresponding author upon reasonable request.

## Acknowledgements

We thank the Nikon Imaging Centre at King’s College London for help with light microscopy. This work was funded by Wellcome Trust Investigator Award 110060/Z/15/Z, BBSRC sLoLa (BB/V003518/1) and a PhD studentship to AP from King’s College London.

## Extended Data

### Supplementary table legends

**Supplementary Table 1.** Lipids detected by LTMS from membrane-associated proteins in this study. The table displays lipid species detected, including length of the acyl chains, molecular formula, m/z, error ppm, retention time (Rt) (min), adduct type, annotation software used for identification and mean value of ion counts with standard deviation (SD) for MyrPalm-GFP (control), Lact-C2-GFP, TOM20-GFP, RACGAP1-GFP, RACGAP1-GFP ΔC1, GFP-CHMP4B or CHMP2A-L-GFP. S stands for samples from cells synchronized at cytokinesis.

**Supplementary Table 2.** Library of lipids detected in untreated wild type HeLa cells and their relative abundance within each lipid family. The table displays the lipid species, mean value of ion counts with standard deviation (SD), and relative abundance within each lipid class in negative or positive modes. According to their physicochemical properties, lipids can be ionized in positive, negative mode, or both (e.g. triacylglycerol (TGs) are detected only in positive mode). These separate experiments are combined in this table for better comparison. Colours indicate lipid species pulled down from RACGAP1-GFP, GFP-CHMP4B or CHMP2A-L-GFP (orange), or Lact-C2 or TOM20-GFP (green).

**Extended Data Fig. 1.**
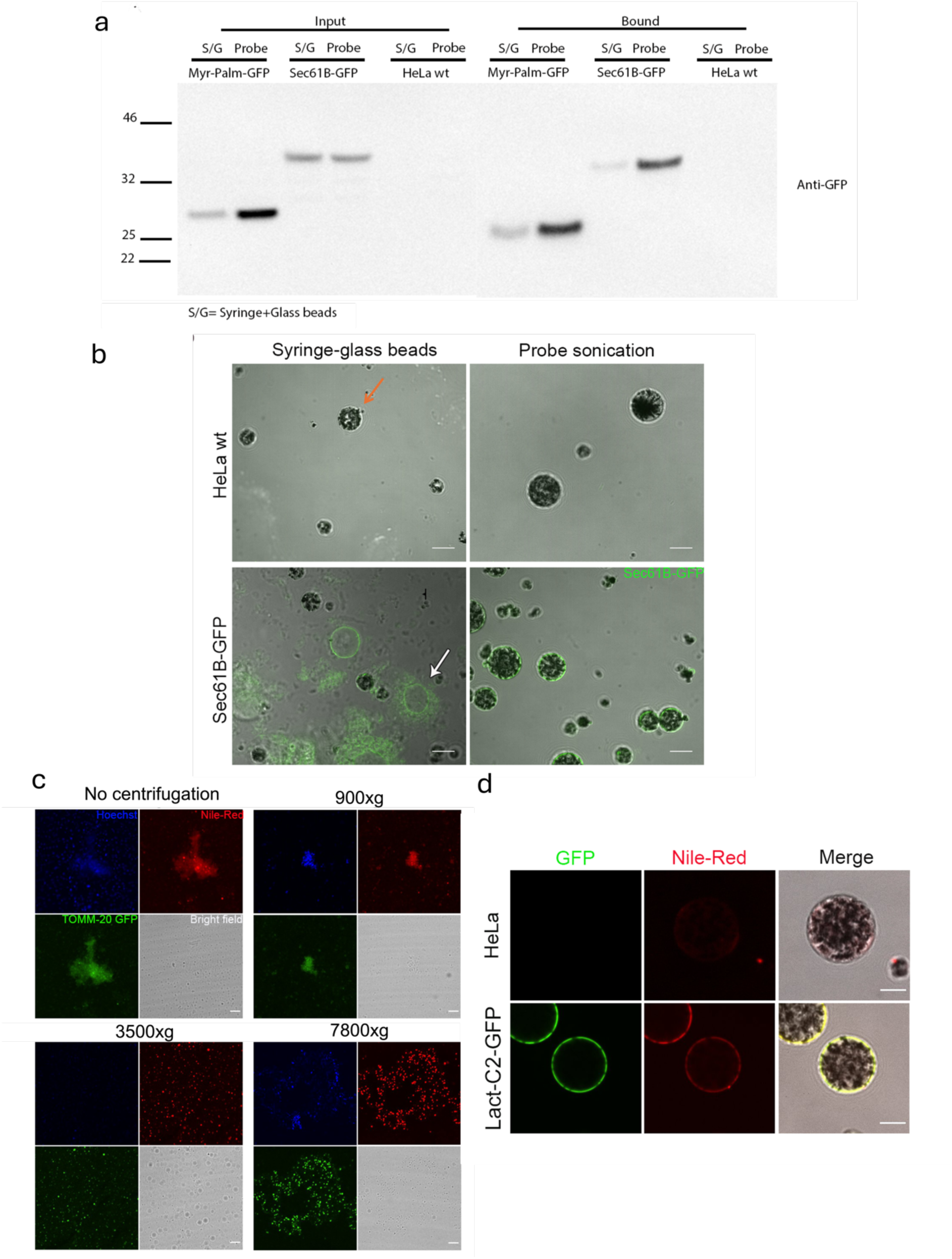
Optimisation of lipid-trap mass spectrometry cell lysis and pulldown conditions. **(a)** Western blot of HeLa wt cells expressing no GFP tag and HeLa cells stably expressing MyrPalm-GFP or SEC61B-GFP, after homogenization with syringe/glass beads (S/G) and probe sonication (Probe), 0.5% of input (left) and 25% of bound fraction to GFP-trap beads (right) were loaded. **(b)** Representative confocal images of GFP-Trap beads after equilibration with cell lysates from HeLa wt or HeLa cells stably expressing SEC61B-GFP (green) after homogenization with syringe/glass beads or probe sonication, scale bar=20μm. Orange arrow: GFP-Trap bead. White arrow: intact ER membrane. **(c)** Confocal images of cell lysate supernatant from HeLa cells stably expressing TOM20-GFP (green) after probe sonication stained with Hoechst (blue, DNA stain) and Nile-Red (red, neutral lipid stain) after different centrifugation speeds (No centrifugation, 900xg, 35000xg, 7800xg), scale bar=10μm. **(d)** Confocal images of GFP-Trap beads after equilibration with cell lysates from HeLa wt and HeLa cells stably expressing Lact-C2-GFP (green) stained with Nile red (red, neutral lipid stain), scale bar=10μm.

**Extended Data Fig. 2.**
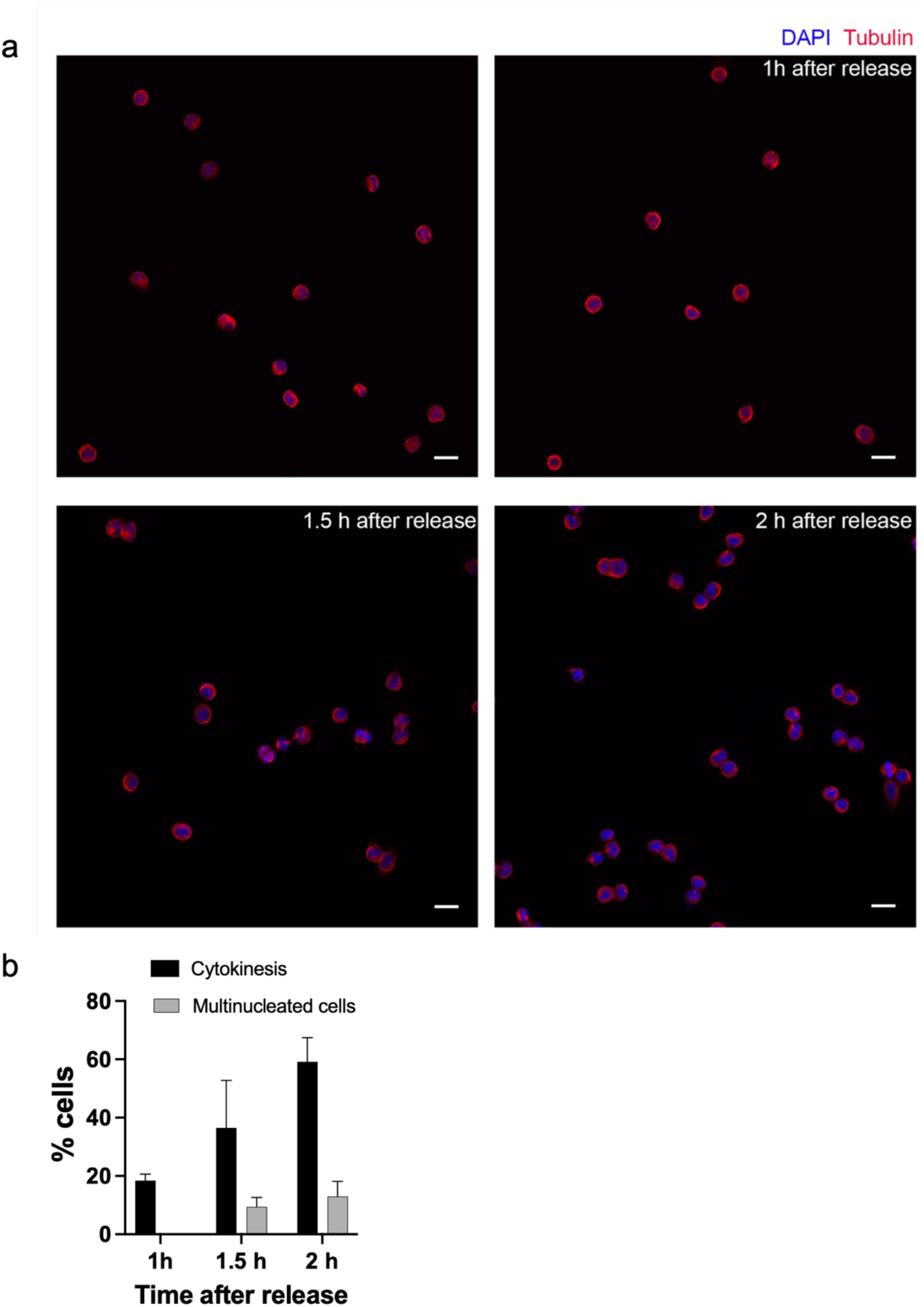
Synchronization of HeLa cells at cytokinesis. **(a)** Representative confocal images of HeLa cells fixed and stained with DAPI (blue) and anti-tubulin antibody (red) to visualize DNA and microtubules. Cells were treated with nocodazole (100 ng/mL) for 12 h, washed and imaged 1h, 1.5h and 2h after mitotic shake-off. Note the highest number of cytokinetic cells is 2h after release. The image was acquired with an A1 confocal inverted microscope using a 40X air lens. Scale bar=20μm. **(b)** Quantification of the percentage of HeLa cells synchronized at cytokinesis at 1h, 1.5h and 2h. Data represent mean value and standard deviation (SD), ~ 60 cells per experiment. N=2 experiments

**Extended Data Fig. 3.**
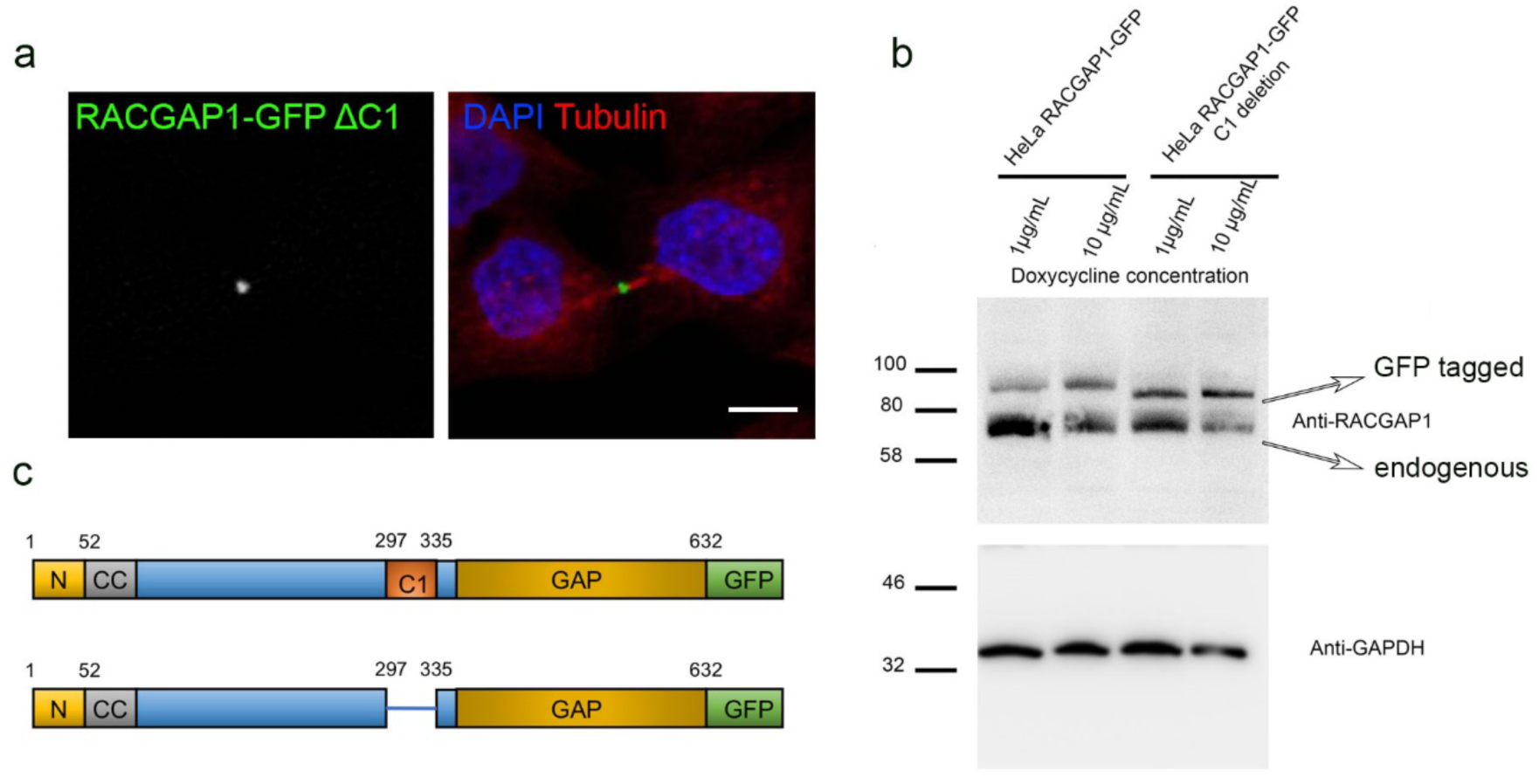
RACGAP1 C1 domain deletion. **(a)** Confocal image of HeLa cell stably expressing RACGAP1-GFP ΔC1 (green) at cytokinesis, fixed and stained with DAPI (blue) and anti-tubulin antibody (red) to visualize DNA and microtubules, respectively scale bar=10μm. **(b)** Western blot of HeLa cells stably expressing RACGAP1 and RACGAP1 ΔC1 showing endogenous RACGAP1 (lower band) and transfected RACGAP1-GFP (upper band, black arrow), GAPDH (loading control). The SDS-PAGE gel was run using the total lysate of HeLa cells stably expressing RACGAP1-GFP and RACGAP1 ΔC1-GFP. **(c)** Cartoon depicting domains of RACGAP1 and its ΔC1 mutant; C1, cysteine-rich domain; CC, coiled-coil; GAP, GTPase-activating domain. Adapted from Lekomtsev *et al.* (*1*).

**Extended Data Fig. 4.**
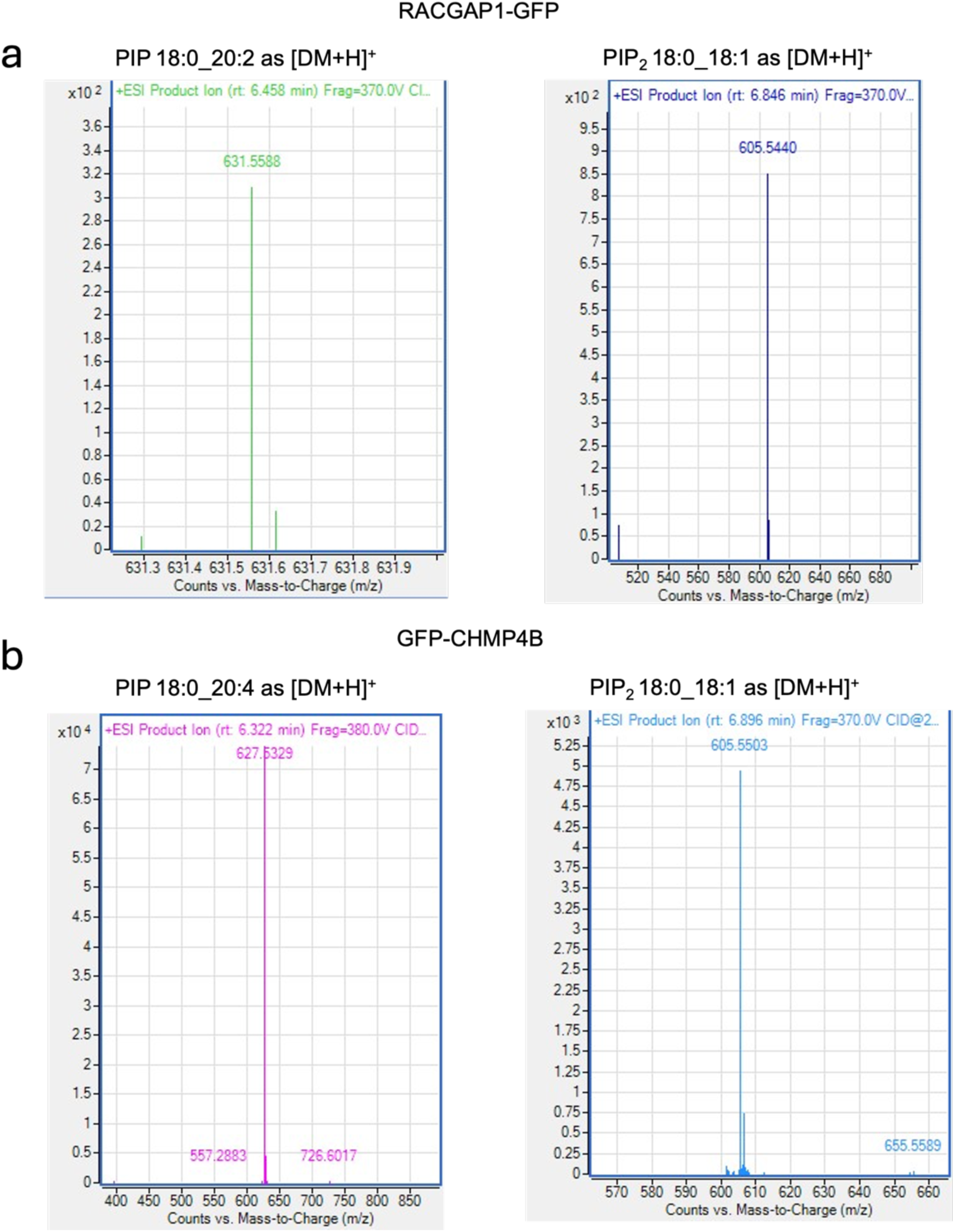
MS2 spectra for derivatized PIPs shown in figures 3b and 4b. **(a)** MS2 spectra acquired for derivatized PIP 18:0_20:2 and PIP_2_ 18:0_18:1 detected from RACGAP1-GFP samples showing diagnostic diacylglycerol (DAG) fragments. **(b)** MS2 spectra acquired for derivatized PIP 18:0_20:4 and PIP_2_ 18:0_18:1 detected from GFP-CHMP4B samples showing diagnostic DAG fragments. MassHunter Qualitative Analysis Software was used to generate the image.

**Extended Data Fig. 5.**
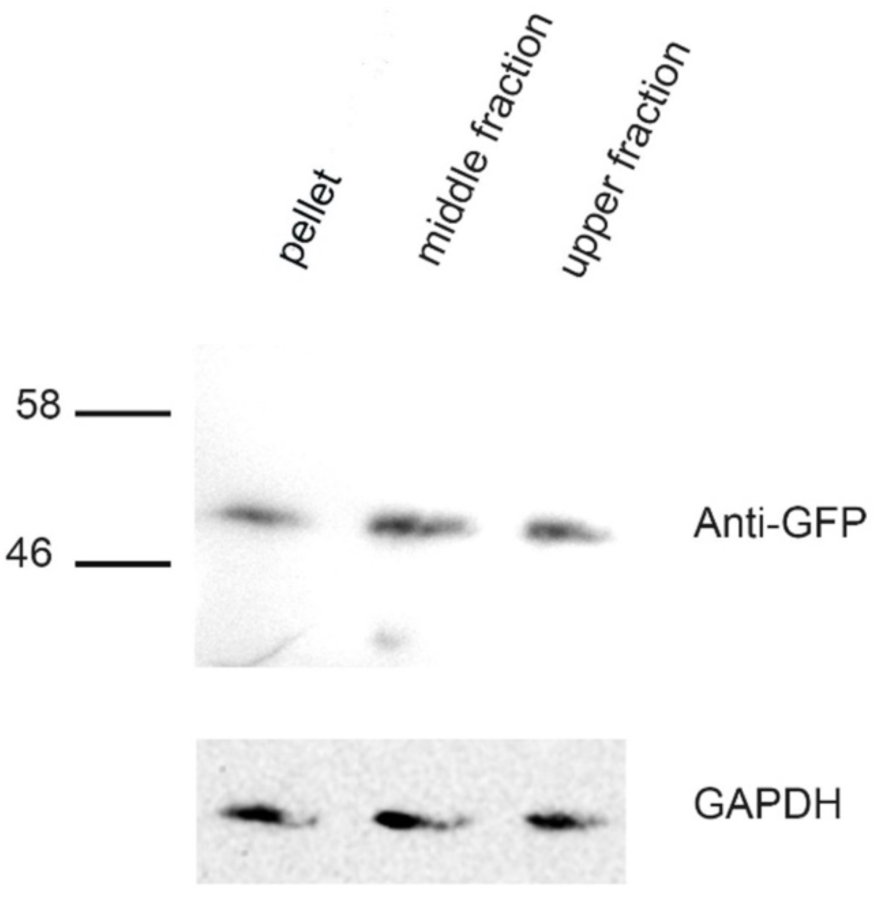
CHMP4B was equally distributed in the fractions and pellet after fractionation by ultracentrifugation. Western blot of HeLa cells stably expressing GFP-CHMP4B showing the protein levels of CHMP4B and GAPDH (loading control) after ultracentrifugation in pellet (plasma and organelle membranes), middle and upper cytoplasmic fractions. The SDS-PAGE gel was run using the total lysate of HeLa cells stably expressing GFP-CHMP4B.

